# Suppression of GABAergic transmission in the spinal dorsal horn induces pain-related behavior in a chicken model of spina bifida

**DOI:** 10.1101/561852

**Authors:** Md. Sakirul I Khan, Hiroaki Nabeka, Farzana Islam, Tetsuya Shimokawa, Shouichiro Saito, Tetsuya Tachibana, Seiji Matsuda

**Affiliations:** Department of Anatomy and Embryology, Graduate School of Medicine, Ehime University, Toon 791-0295, Ehime, Japan; Laboratory of Veterinary Anatomy, Faculty of Applied Biological Sciences, Gifu University, Yanagido, 501-1193, Gifu, Japan; Department of Agrobiological Science, Faculty of Agriculture, Ehime University, Matsuyama 790-8566, Japan

**Keywords:** Spina bifida, Chicken model, Pain-related behavior, GABAergic transmission, Apoptosis

## Abstract

Spina bifida aperta (SBA), one of the most common congenital malformations, causes various neurological disorders. Pain is a common complaint of patients with SBA. However, little is known about the neuropathology of SBA-related pain. Because loss of γ-aminobutyric acid (GABA)ergic neurons in the spinal cord dorsal horn is associated with pain, we hypothesized the existence of cross-talk between SBA-related pain and alterations in GABAergic transmission in the spinal cord. Therefore, we investigated the kinetics of GABAergic transmission in the spinal cord dorsal horn in a chicken model of SBA. Neonatal chicks with SBA exhibited various pain-like behaviors, such as an increased number of vocalizations with elevated intensity (loudness) and frequency (pitch), reduced mobility, difficulty with locomotion, and escape reactions. Furthermore, the chicks with SBA did not respond to standard toe-pinching, indicating disruption of the spinal cord sensorimotor networks. These behavioral observations were concomitant with loss of GABAergic transmission in the spinal cord dorsal horn. We also found apoptosis of GABAergic neurons in the superficial dorsal horn in the early neonatal period, although cellular abnormalization and propagation of neurodegenerative signals were evident at middle to advanced gestational stages. In conclusion, ablation of GABAergic neurons induced alterations in spinal cord neuronal networks, providing novel insights into the pathophysiology of SBA-related pain-like complications.

## Introduction

Spina bifida aperta (SBA), a neural-tube defect (NTD) that causes lifelong neurological complications, develops in approximately 1 in 1,000 neonates worldwide [1]. SBA is primarily characterized by defective fusion of the neural tube, which causes *in utero* deformities in the exposed spinal [2–5]. Such spinal cord deformities lead to varying degrees of motor and/or sensory deficits, resulting in neurological disorders such as spinal ataxia, paralysis of the legs, problems with bowel and bladder control, and pain complications [5–8]. NTD-induced pain is a common complaint of SBA patients of any age [8–10]. Individuals with SBA frequently have various risk factors for pain, such as musculoskeletal deformities, clogged/infected shunts, urinary tract infections, bowel problems, and suboptimal positioning [11,12]. However, little is known about the pathophysiology of pain in SBA. Thus, research in animal models of SBA is needed to better understand the underlying cellular and molecular mechanisms and to develop novel therapeutic interventions.

The pain experienced by people with NTD may be nociceptive or neuropathic or a combination thereof [8]. A loss of inhibitory transmission in the spinal cord dorsal horn is important in the development of chronic pain. Spinal cord injury-induced loss of γ-aminobutyric acid (GABA)ergic neurons, the principle inhibitory interneurons, in the superficial dorsal horn causes persistent pain [13–18]. These facts led us to postulate the existence of cross-talk between SBA-related pain complications and alteration in GABAergic transmission in the spinal cord dorsal horn. Accordingly, we explored the kinetics of GABAergic transmission in the spinal cord dorsal horn from gestation to post-hatching in a chicken model of SBA to obtain pathophysiological data on the sequence of events associated with SBA-like neurological disorders [19–21].

## Materials and Methods

### Animals

The fertilized eggs of chickens (*Gallus gallus*; Mori Hatchery, Kagawa, Japan) were incubated in a commercial incubator (Showa Furanki, Saitama, Japan) at 37.8 ± 0.2°C with 60% relative humidity to obtain embryos at developmental stages 17–21 inclusive on gestational days 2.5–3.5. The developmental stage of each embryo was determined using the developmental table of Hamburger and Hamilton [22]. The embryos were divided into two groups: the SBA group, in which the roof plate of the neural tube was incised; and the normal control group, in which the neural tube was left intact. The hatched chicks were raised in a room maintained at 30°C with continuous illumination and were fed a commercial diet (crude protein, 24%; metabolizable energy, 3,050 kcal/kg; Toyohashi Feed Mills Co. Ltd., Toyohashi, Japan) with water available *ad libitum*. Because neonatal SBA chicks have difficulty consuming food, they were gavaged a feed slurry (tube feeding) at a mass of 4.0% of their body weight into the crop four times daily. The feed slurry was made by mixing 40% powdered diet with 60% distilled water on a weight basis. The animal experimental protocols were approved by the Committee on the Ethics of Animal Experiments of the Ehime University Graduate School of Medicine, Toon, Japan (No. 05A27-10).

### Surgical manipulation to generate SBA chicks

To produce SBA chicks, surgical manipulation of the neural plate was carried out as described previously [19, 21]. Briefly, the eggshell and amnion were opened and placed under a stereomicroscope to determine the developmental stage of the embryo. Next, the roof plate of the neural tube was incised longitudinally, starting at the level of the cranial margin of the 26th somite, which forms the sixth and seventh thoracic segments, using a custom-made microknife [23, 24]. The incision extended caudally for a distance equivalent to the length of seven somites and was made by inserting the microknife into the neural tube to approximately half the depth of the tube. The roof plate was incised, and care was taken not to damage other parts of the neural tube. After the incision was made and the surgical manipulation was complete, the shell window was closed using transparent adhesive tape and the eggs were re-incubated at 37.8 ± 0.2°C with 60% relative humidity.

### Behavioral assessments

For behavioral assessments, the SBA and control chicks used at postnatal day (PD)-0 were also used at PD-2, PD-4, and PD-10. To characterize pain-related behavioral changes, the spontaneous activities of the normal and SBA chicks were videotaped at 08:00 and 20:00 for 30 min at PD-0, PD-2, PD-4, and PD-10 and scored offline. The intensity and number of vocalizations, the primary indicators of pain in birds [25, 26] were characterized by two independent observers. The intensity of vocalization (Frequency Analyzer 2.0; free software), and number of vocalizations per 10 min were assessed in the SBA and normal chicks.

To elicit a sensorimotor reaction, forceps were used to conduct a standard pinch test of the toes on both legs of the normal and SBA chicks; the tests were repeated three times per chick. The results were considered conclusive and a normal reaction was recorded only if an identical response was obtained on at least three toes in each leg in three consecutive tests. The reactions of the chicks were videotaped and the leg withdrawal reflex, jumping, and vocalization behaviors were scored offline by two independent observers. Behavior was analyzed in six chicks per group at each age.

### Sample collection and tissue preparation

Spinal cord sections from the open defect area (lumbosacral region) from SBA chicks, and from a similar location in the normal chicks, were collected on embryonic day (ED)-14, ED-18, PD-2, PD-4, and PD-10. The incisions of the roof plates of the neural tubes of SBA embryos were confirmed by observing open defects on the backs of the chicks, and the induction of SBA in the chicks post-hatching was confirmed by observing open defects on the backs of the chicks with leg dysfunction ([21]; Fig. 1a; Suppl. Video 1). Six chicks from each group at each age were transcardially perfused with a fixative solution containing 4% paraformaldehyde with 0.5% glutaraldehyde in 0.1 M phosphate-buffered saline (PBS). Next, the spinal cord was removed from the location of the open defect (exposed area, lesion location). Spinal tissue samples from the normal control chicks were collected from locations similar to the tissue collection locations of SBA chicks. The collected tissues were immersed in 4% paraformaldehyde overnight at 4°C, dehydrated, and embedded in paraffin.

**Figure 1.**
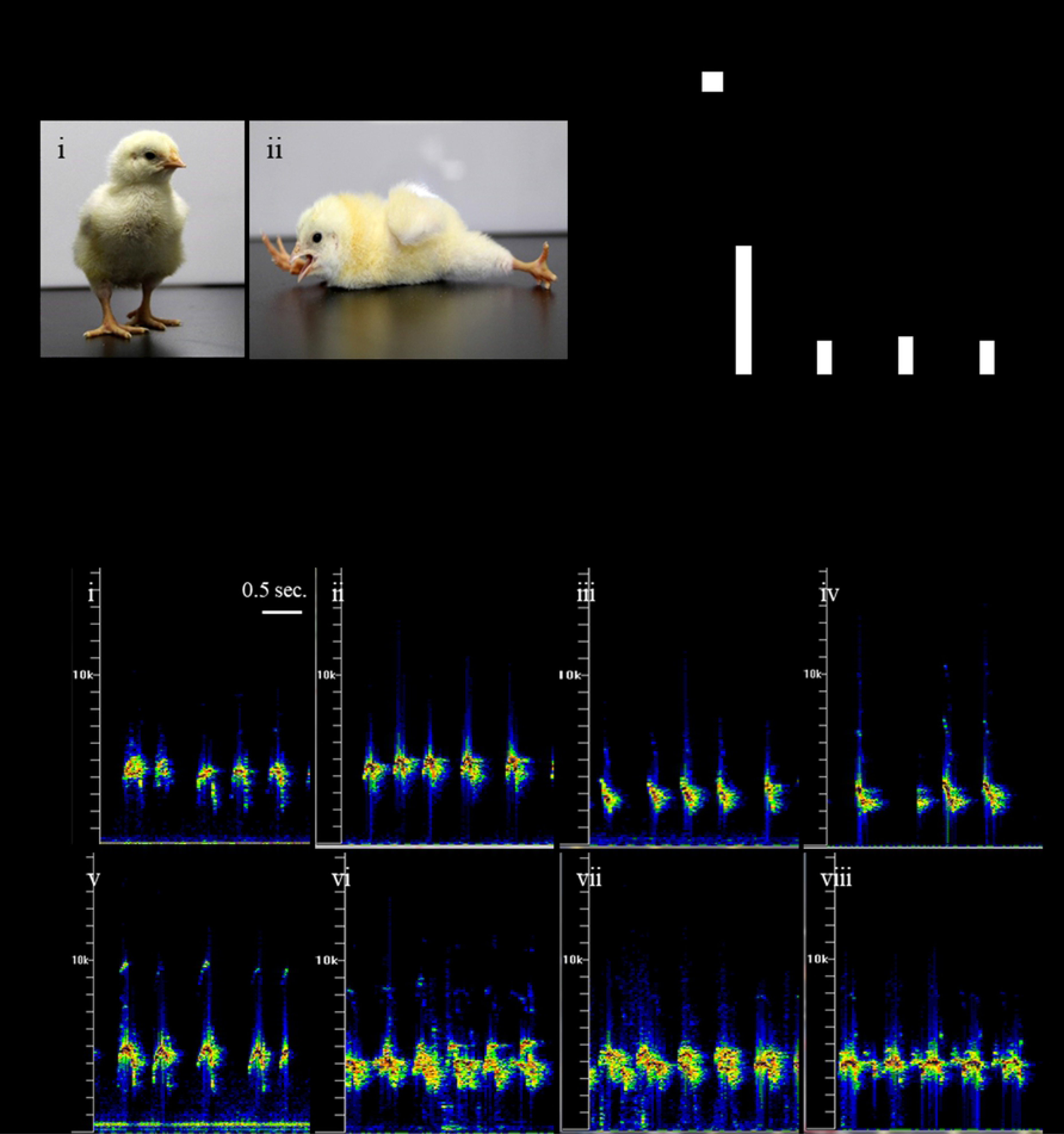
Pain-like behavioral changes in spina bifida aperta (SBA) chicks. (a) Representative photographs of the phenotypes of normal and SBA chicks. The control chicks were able to move freely (i) whereas SBA chicks moved with severe difficulty (ii). (b) Numbers of vocalizations in normal and SBA neonatal chicks. (c) The intensity (loudness) and frequency (pitch) of vocalizations in normal and SBA neonatal chicks. The data in Fig. 1b are means ± SEM; n = 6 per group at each age. *Significantly different from the SBA group (*P* < 0.001), two-way ANOVA and *post hoc* Tukey’s test.

### Immunohistochemical staining

The avidin-biotin complex method was used to analyze GABAergic immunoreactivities. Briefly, spinal cord sections (7 µm) from the location of the open defect (exposed area) of the lumbosacral cord in the SBA chicks, and from a similar location in the normal chicks, were deparaffinized, rehydrated, and treated with PBS containing 10% methanol and 3% hydrogen peroxide (H_2_O_2_) for 10 min. After rinsing with PBS, the spinal cord sections were treated with 5% bovine serum albumin, 1% normal goat serum, 0.1% fish gelatin, and 0.1% NaN_3_ in PBS for 1 h followed by incubation with the primary antibodies, rabbit polyclonal anti-GABA (1:1,500; Sigma, St. Louis, MO) overnight at 4°C, or mouse monoclonal anti-parvalbumin (PV) (1:500; Sigma), mouse monoclonal anti-calbindin-D-28K (CB), or mouse monoclonal anti-calretinin (CR) (1:500; SWANT, Bellinzona, Switzerland) for 60 h at 4°C. Next, the sections were washed three times for 10 min each in PBS and reacted with the secondary antibodies for 2 h at room temperature. After rinsing three times for 10 min each in PBS, the avidin-biotin-peroxidase complex (1:300; Dako, Glostrup, Denmark) was applied for 1 h at room temperature. The sections were immersed in 3,3-diaminobenzidine (DAB) (Sigma) containing 0.0033% H_2_O_2_ for approximately 10 min. After rinsing with distilled water, the sections were dehydrated, mounted, and visualized under a light microscope (Nikon, Tokyo, Japan).

Antibody specificity was tested using a negative staining procedure with normal rabbit IgG (1:250; Dako, Glostrup, Denmark) instead of the primary antibodies; the samples were processed as described above. There was a lack of non-specific staining (data not shown).

### Quantitative analysis of GABAergic neurons

For quantitative analysis of GABA-expressing neurons, serial coronal sections (7 µm) from the location of the open defect (exposed area) of the lumbar cord in the SBA chicks, and from a similar location in the normal chicks, were stained with a rabbit polyclonal anti-GABA antibody (1:1,500; Sigma). The tissues were observed under a Nikon Eclipse E800 light microscope, and images were acquired using a charge-coupled device (CCD) camera attached to the microscope (Nikon Digital Sight DS-L2). For analysis of GABA-expressing neurons, all GABA-immunopositive cells with a rounded profile in superficial dorsal horn laminae I–III were considered and counted. The total number of neurons in the superficial dorsal horn on each side of the spinal cord was calculated and averaged across six cross sections per chick at each age; each group comprised six chicks of each age.

For quantitative analysis of PV-, CB-, and CR-expressing neurons, serial coronal sections were stained with mouse monoclonal anti-PV (1:500; Sigma), mouse monoclonal anti-CB, or mouse monoclonal anti-CR (1:500; SWANT) antibody, and PV-, CB-, and CR-immunopositive cells were enumerated as described above.

### Measurement of staining intensity

The GABAergic immunoreactivities in the dorsal horn lamina I–III of the open defect in the spinal cord of SBA chicks, and at a similar location in normal chicks, were measured using ImageJ software (National Institutes of Health, Bethesda, MD, USA). To correct for background, three background intensity readings were taken per image. These readings were averaged and subtracted from the signal intensity to yield the protein staining intensity. The intensity data are presented as the mean ± SEM. Six random sections per chick at each age in each group were analyzed for quantification; each group comprised six chicks of each age.

### Immunofluorescence staining

Immunofluorescence staining was performed as described previously [21]. For double immunofluorescence staining, the sections were incubated for 60 h at 4°C in a solution containing rabbit polyclonal anti-GABA (1:3,000; Sigma) plus mouse monoclonal anti-CB or mouse monoclonal anti-CR (1:1000; SWANT) antibodies; and rabbit polyclonal anti-caspase 3 (1:1000; Bioss) plus mouse monoclonal anti-CB (1:1000; SWANT) antibodies. After washing in PBS, the sections were treated for 2 h at room temperature with an Alexa Fluor 546-conjugated goat anti-rabbit IgG (H+L) (1:1,000; Invitrogen) or an Alexa Fluor 488-conjugated goat anti-mouse IgG (H+L) (1:1,000; Invitrogen) and 4′,6-diamidino-2-phenylindole (DAPI), washed with PBS, mounted with Vectashield (Vector Laboratories), and visualized using a Nikon A1 confocal microscope equipped with a 100× objective lens (Nikon).

### Statistical analysis

Statistical analysis was performed on the mean values and data are reported as the mean ± SEM. The data were subjected to two-way analysis of variance (ANOVA) with the Tukey–Kramer *post hoc* test, and *P*-values < 0.05 were considered to indicate statistical significance.

## Results

### Assessment of behavioral disorders

Various neurological complications were evident in neonatal chicks with SBA (Suppl. Video 1). Difficulty during locomotion (Fig. 1a) and increased numbers of vocalizations (Fig. 1b), with greater intensity (loudness) and frequency (pitch; Fig. 1c), were observed in SBA chicks compared with control chicks during the early neonatal period. Chicks with SBA also showed reduced confidence in mobility, escape reactions, anxiety, fear or restlessness, and inappetence. These behavioral changes are considered primary indicators of pain in birds [25, 26], indicating that the chicks with SBA experienced pain.

The sensorimotor reactions of the chicks are summarized in Table 1. In normal chicks, toe-pinching resulted in withdrawal of the limb, jumping, and vocalization (Supplementary Video 2). The responses were consistent in control chicks at PD-0, PD-2, PD-4, and PD-10; therefore, the pattern was considered a normal sensorimotor reaction. In the first few hours after hatching (PD-0), toe-pinching elicited relatively normal reactions in SBA chicks compared to age-matched control chicks. However, commencing on PD-2, the sensorimotor reactions gradually decreased; there was almost no response at PD-10 (Table 1; Supplementary Video 1), indicating disruption of the sensorimotor networks in the chicks with SBA.

**Table 1.** Assessment of sensorimotor reactions in spina bifida aperta (SBA) chicks. A 5-point scoring system was used to grade sensorimotor reactions (leg withdrawal reflex, jumping, and vocalizations) after toe-pinching in normal control and SBA chicks at the early neonatal stage. Sensorimotor reactions were graded as no reaction (0 points), minimal reaction (1–2 points), reduced reaction (3–4 points), and normal reaction (5 points). * Difficult to distinguish toe-pinching-induced vocalization reaction, as the SBA chicks were experienced persistent vocalization.

| Age | Sensorimotor reactions by toe-pinching (total score: 5 per parameter) |  |  |  |  |  |
| --- | --- | --- | --- | --- | --- | --- |
|  | Control |  |  | SBA |  |  |
|  | Leg withdrawal | Vocalization | Jump | Leg withdrawal | Vocalization | Jump |
| PD-0 | 4 | 4 | 5 | 5 | 4 | 5 |
| PD-2 | 5 | 5 | 5 | 3 | 5 | 2 |
| PD-4 | 5 | 5 | 5 | 2 | 5* | 1 |
| PD-10 | 4 | 3 | 4 | 1 | 5* | 0 |

### Expression of GABA in the dorsal horn of the spinal cord

The GABA immunoreactivity in exposed spinal cord in embryonic and neonatal chicks is shown in Fig. 2. GABA immunoreactivity was detected in the spinal cord, predominantly in dorsal horn laminae I–III. Such GABA immunoreactivity in avian species is in line with mammalian studies of GABA distribution in the spinal cord [27]. GABA immunoreactivity was higher in SBA chicks than in normal chicks at the embryonic stage (ED-14). In contrast, GABA immunoreactivity was lower in SBA chicks at the neonatal stage (PD-4) compared to not only age-matched normal chicks but also SBA chicks at the embryonic stage. In fact, at ED-14, GABA expression was significantly higher in SBA chicks than in age-matched control chicks, whereas the opposite pattern was evident at PD-4 (Fig. 2i). At ED-14, the number of GABA-immunopositive cells was significantly higher in dorsal horn laminae I–III of SBA chicks than in the control chicks, but the number decreased at PD-4 (Fig. 2j).

**Figure 2.**
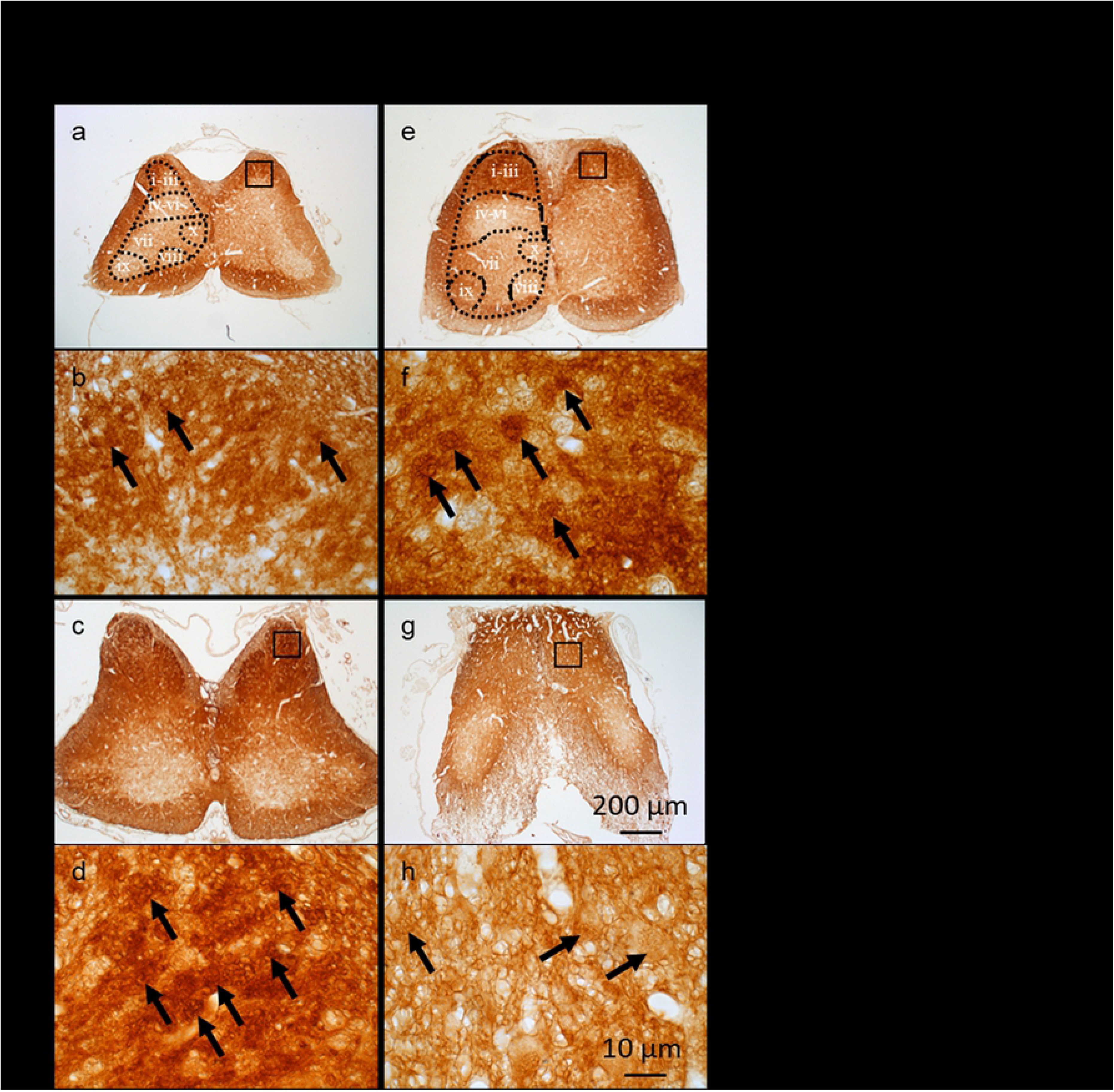
Postnatal loss of γ-aminobutyric acid (GABA) immunoreactivity in the spinal cord dorsal horns of SBA chicks. (a–h) Immunohistochemical analysis of the expression and localization of GABA in the spinal cord open defect area in SBA chicks, and at a similar location in normal chicks, at ED-14 and PD-4. Representative images of the paraffin-embedded spinal cords of control (a–d) and SBA (e–h) chicks. (i) GABA immunoreactivity in the dorsal horn laminae I–III of control and SBA chicks measured using ImageJ software. The optical density (OD) was calculated using the following formula: OD = (log_10_ [incident light/transmitted light]). Six random sections per chick at each age and for each group were analyzed. (j) Number of GABA-immunopositive cells in the spinal cord dorsal horn laminae I–III of control and SBA chicks. The data in Figs. 2i and 2j are means ± SEM; n = 6 in each group at each age. *Significantly different from the SBA group at each age (*P* < 0.01), two-way ANOVA and *post hoc* Tukey’s test. Arrows indicate GABA-immunopositive cells.

### Expression of calcium-binding GABAergic subpopulations in the spinal cord dorsal horn

Figures 3, 4, and 5 show the immunoreactivities of the PV, CR, and CB calcium-binding GABAergic subpopulations, respectively. PV-positive neurons were predominantly detected in the ventral horn area, but also in the dorsal horn (Fig. 3). The overall density of PV-immunopositive fibers in the spinal cord increased with increasing developmental stage in the normal chicks (Fig. 3a–d) but not in the SBA chicks (Fig. 3e–h). Similarly, the number of PV-positive neurons in dorsal horn laminae I–III at ED-14 was greater in SBA chicks than in normal chicks; however, after hatching (at P4), the opposite pattern was evident (Fig. 3j). PV immunoreactivity at ED-14 was significantly greater in the SBA chicks than in age-matched control chicks; the opposite pattern was evident at the neonatal stage (Fig. 3i).

**Figure 3.**
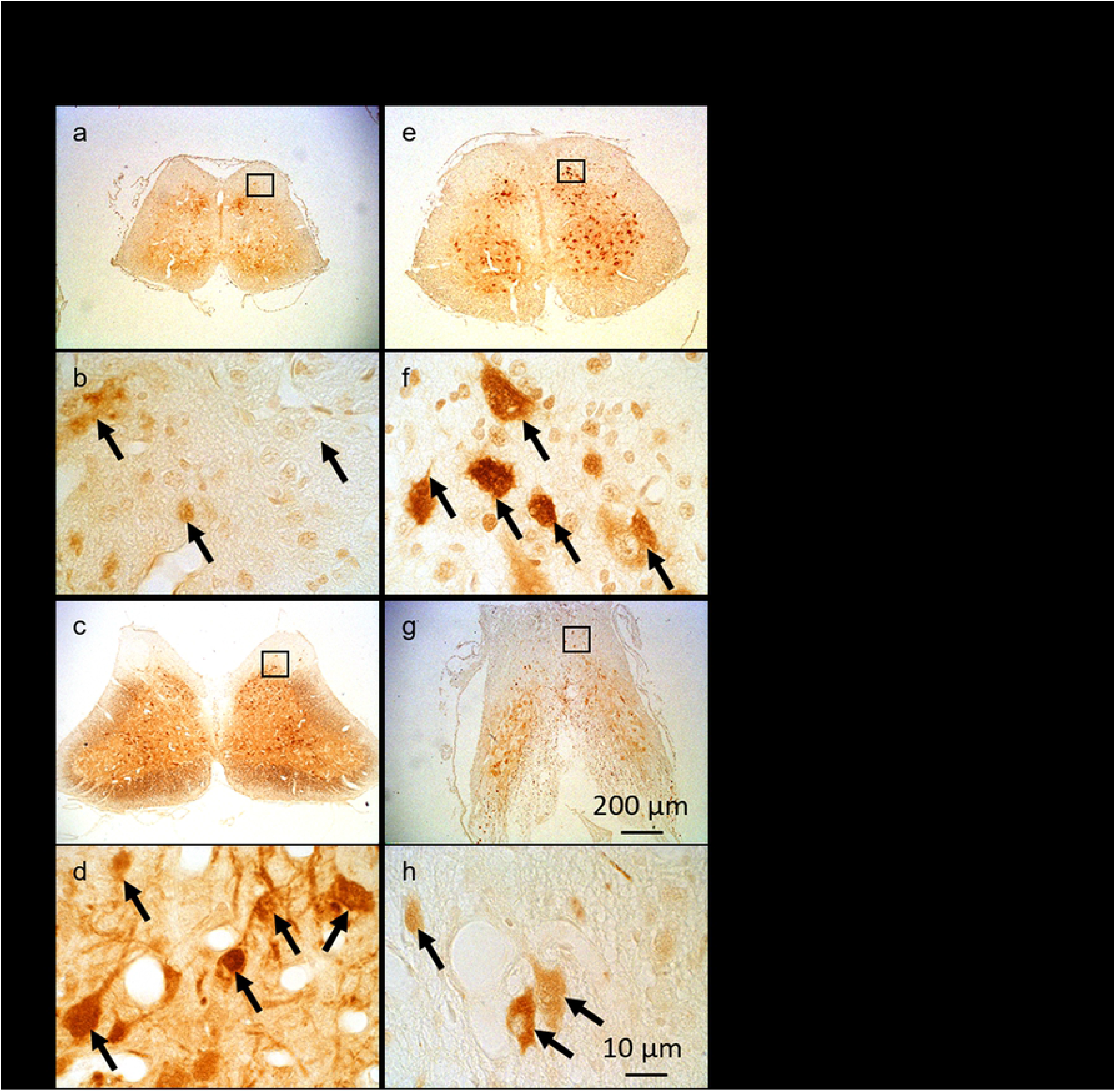
Postnatal loss of parvalbumin (PV) immunoreactivity in the spinal cord dorsal horns of SBA chicks. (a–h) Immunohistochemical analyses of the expression and localization of PV in the spinal cord open defect area in SBA chicks, and at a similar location in normal chicks, at ED-14 and PD-4. Representative images of the paraffin-embedded spinal cords of control (a–d) and SBA (e–h) chicks. (i) PV immunoreactivity in the dorsal horn laminae I–III of control and SBA chicks measured using ImageJ software. The optical density (OD) was calculated using the following formula: OD = (log_10_ [incident light/transmitted light]). Six random sections per chick at each age and in each group were analyzed. (j) Numbers of PV-immunopositive cells in the spinal cord dorsal horn laminae I–III of control and SBA chicks. The data in Figs. 2i and 2j are presented as means ± SEM; n = 6 per group at each age. *Significantly different from the SBA group at each age (*P* < 0.01), two-way ANOVA and *post hoc* Tukey’s test. Arrows indicate PV-immunopositive cells.

**Figure 4.**
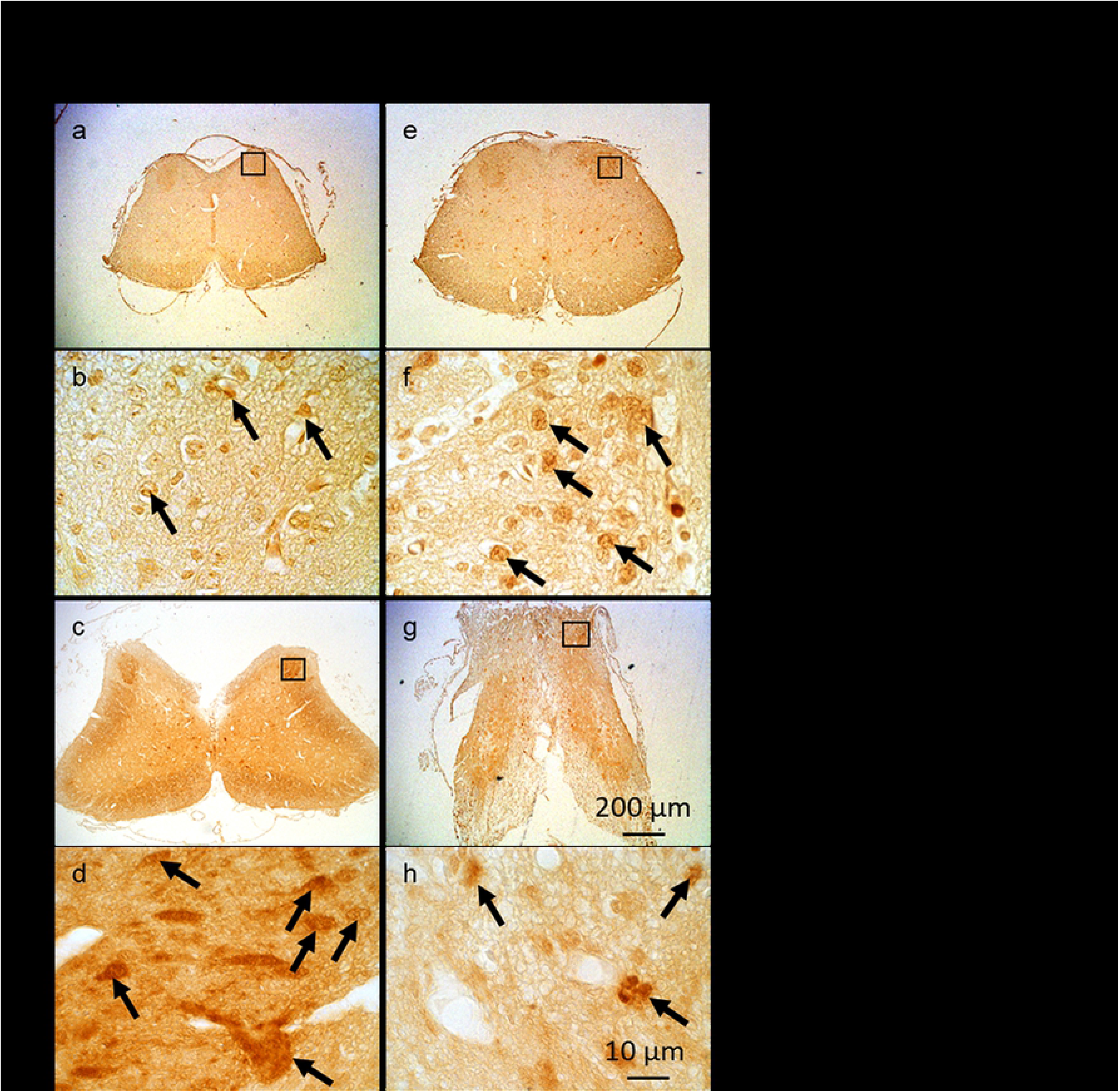
Postnatal loss of calretinin (CR) immunoreactivity in the spinal cord dorsal horns of SBA chicks. (a–h) Immunohistochemical analyses of the expression and localization of CR in the spinal cord open defect area in SBA chicks, and at a similar location in normal chicks, at ED-14 and PD-4. Representative images of the paraffin-embedded spinal cords of control (a–d) and SBA (e–h) chicks. (i) CR immunoreactivity in the dorsal horn laminae I–III of control and SBA chicks measured using ImageJ software. The optical density (OD) was calculated using the following formula: OD = (log_10_ [incident light/transmitted light]). Six random sections per chick at each age and for each group were analyzed. (j) Numbers of CR-immunopositive cells in the spinal cord dorsal horn laminae I–III of control and SBA chicks. The data in Figs. 2i and 2j are means ± SEM; n = 6 per group at each age. *Significantly different from the SBA group at each age (*P* < 0.01), two-way ANOVA and *post hoc* Tukey’s test. Arrows indicate CR-immunopositive cells.

**Figure 5.**
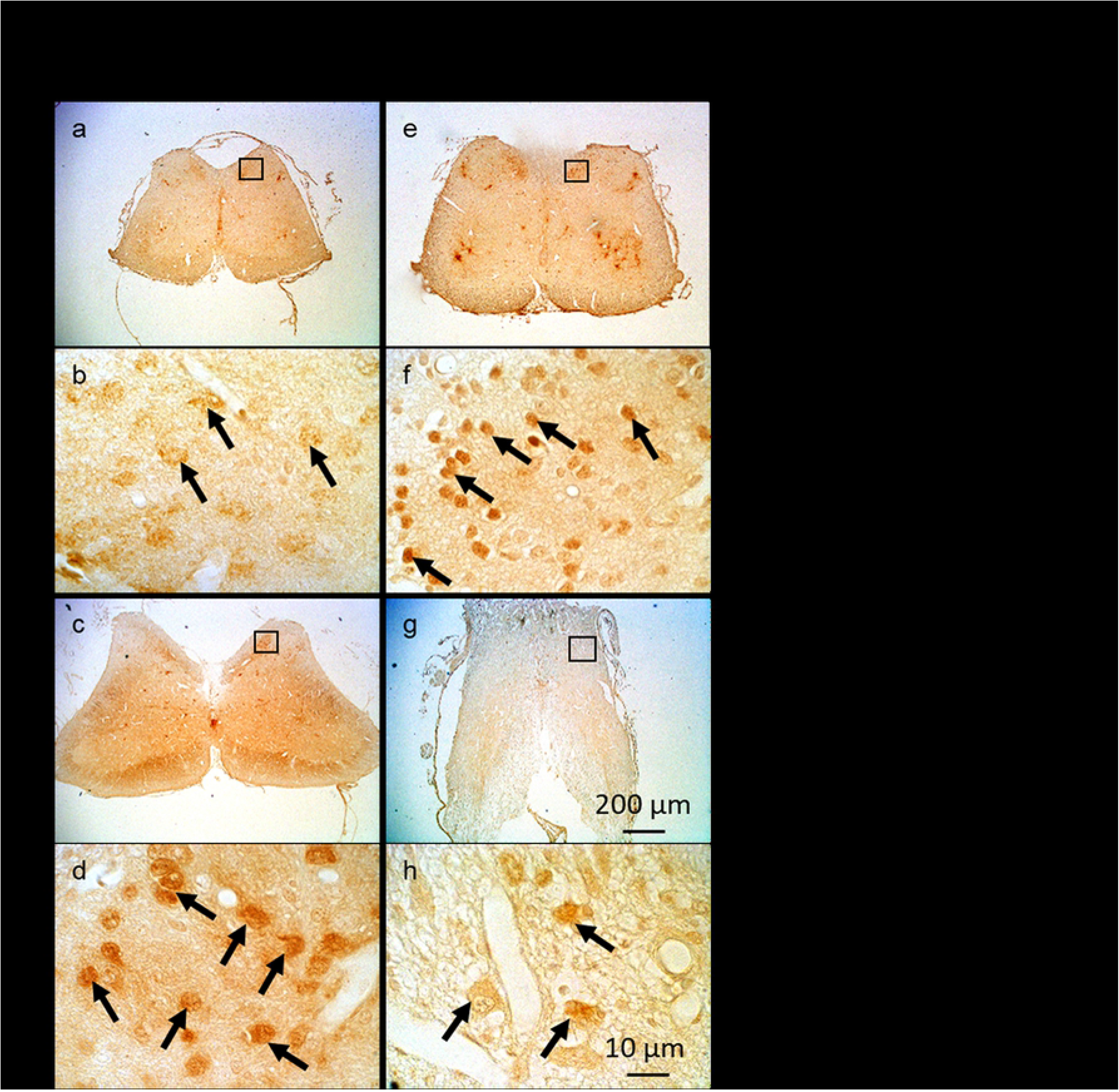
Postnatal loss of calbindin-D-28K (CB) immunoreactivity in the spinal cord dorsal horns of SBA chicks. (a–h) Immunohistochemical analyses of the expression and localization of 22 CB in the spinal cord open defect area in SBA chicks, and at a similar location in normal chicks, at ED14 and PD-4. Representative images of the paraffin-embedded spinal cords of control (a–d) and SBA (e–h) chicks. (i) CB immunoreactivity in the dorsal horn laminae I–III of control and SBA chicks measured using ImageJ software. The optical density (OD) was calculated using the following formula: OD = (log_10_ [incident light/transmitted light]). Six random sections per chick at each age and for each group were analyzed. (j) Numbers of CB-immunopositive cells in the spinal cord dorsal horn laminae I–III of control and SBA chicks. The data in Figs. 2i and 2j are presented as means ± SEM; n = 6 per group at each age. *Significantly different from the SBA group at each age (*P* < 0.01), two-way ANOVA and *post hoc* Tukey’s test. Arrows indicate CB-immunopositive cells.

CR immunoreactivity in the spinal cord of the SBA and normal chicks is shown in Fig. 4. A few intensely labeled CR-immunopositive cells were observed in both the normal and SBA chicks at the embryonic stage, and this was relatively prominent in dorsal horn laminae I–III of the SBA chicks. Although the CR immunoreactivity of cells and fibers increased in the spinal cords of neonatal normal chicks (Fig. 4a–d), CR immunoreactivity was lower in SBA chicks at PD-4 (Fig. 4e–h). Additionally, the number and staining intensity (Fig. 4i, j) of CR-expressing neurons in dorsal horn laminae I–III at PD-4 were significantly reduced in SBA chicks compared to the control chicks.

CB immunoreactivity in the spinal cords of SBA and normal chicks is shown in Figure 5. CB-immunopositive cells were detected in the dorsal and ventral horns of the spinal cord. In the ventral horn, strongly CB-immunoreactive small neurons were distributed in lamina VII of both the normal and SBA chicks; in mammals, these are reportedly Renshaw cells [28, 29]. The number and staining intensity of CB-immunopositive cell bodies and fibers differed between the control and SBA chicks (Fig. 5i, j). In the SBA chicks, the number of CB-immunopositive cells in laminae I–III during the embryonic stage was higher than in normal chicks. However, post-hatching, the number and immunoreactivities of CB-immunopositive neurons were lower in SBA chicks than in normal chicks.

### GABAergic transmission in the dorsal horn of the spinal cord

To determine whether GABAergic neurons were colocalized in spinal cord dorsal horn and the pain-like behavior exhibited by SBA chicks was associated with alterations in GABAergic transmission, double immunofluorescence staining of GABA and CB or CR in dorsal horn laminae I–III of the spinal cord was performed at ED-14 to PD-10. Double staining of GABA and CB (Fig. 6a, b, e, f) or CR (Fig. 6i, j, m, n) increased during the later stages of the embryonic period. This staining was very strong at ED-18 (Fig. 6f, n), but decreased after PD-4 (Fig. 6g, h, o, p), the time point at which the SBA chicks showed severe pain-like behavioral changes (Fig. 1, Suppl. Video 1).

**Figure 6.**
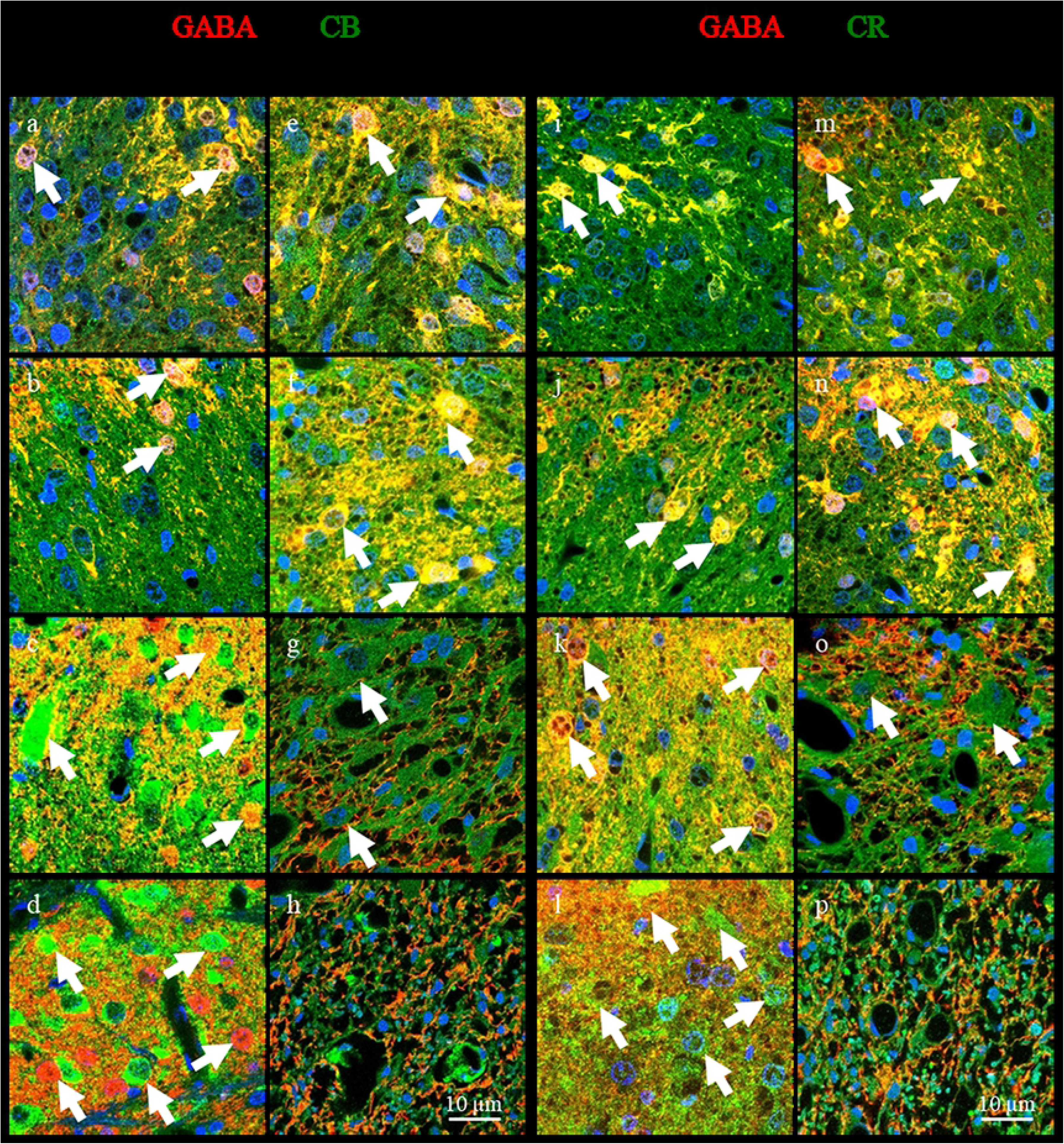
Postnatal loss of GABAergic transmission in the spinal cord dorsal horns of SBA chicks. (a) Double immunofluorescence staining for GABA and either CB or CR. Representative confocal images of the localization of GABA and CB (a–h) or GABA and CR (i–p) in normal and SBA chicks on ED-14, ED-18, PD-4, and PD-10. Images of the dorsal horn (laminae I–III) in the open defect of the lumbar cord in SBA chicks and in a similar location in normal chicks. GABA, red; CB or CR, green; co-localization, yellow; DAPI, blue. Arrows indicate GABAergic-immunopositive cells.

### Neurodegeneration in the spinal cord

To clarify the mechanisms underlying the loss of GABAergic neurons in SBA chicks, the immunoreactivity of caspase-3, a marker of apoptosis, in CB-expressing neurons was evaluated in the dorsal horn laminae I–III of the spinal cord (Fig. 7). In SBA chicks, weak caspase-3 immunoreactivity was evident in CB-expressing neurons at ED-18 (Fig. 7v), and moderate to high caspase-3 immunoreactivity was found at PD-2 and PD-4 (Fig. 7vi and vii). However, most of these GABAergic neurons in the dorsal horn had likely degenerated by PD-10, because only a few interneuron-like structures remained at this point (Fig. 7viii), suggesting that the degeneration of GABAergic neurons in the dorsal horn of the exposed cord of the SBA chicks was due to apoptosis. In addition to the GABAergic neurons, strong caspase-3 immunoreactivity was observed in the tissue of the exposed cord in the SBA chicks at P4 (Fig. 7vii), suggesting that the loss of tissue area may also be due to apoptosis.

**Figure 7.**
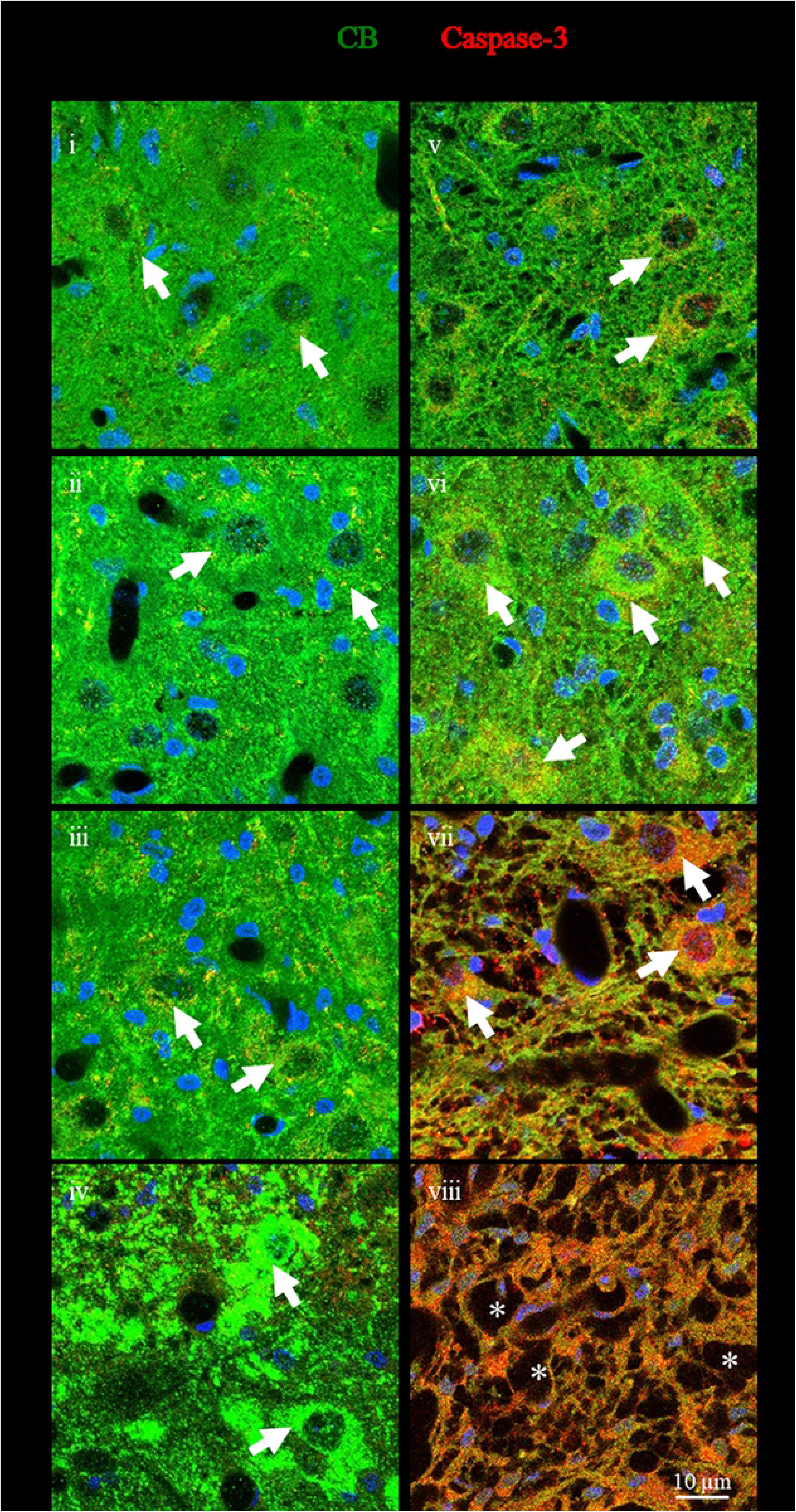
Caspase 3-mediated apoptosis of GABAergic neurons in the spinal cord dorsal horns of SBA chicks. (i–viii) Representative confocal images showing immunofluorescence staining for caspase-3, a marker of apoptosis, and CB-expressing neurons, in the dorsal horn open defect in the lumbar cord of SBA chicks (v–viii) and at a similar location in normal chicks (i–iv). Caspase-3, red; CB, green; co-localization, yellow; DAPI, blue. *Interneuron-like damaged structures in SBA chicks at PD-10. Arrows indicate caspase-3 immunoreactivity in CB-expressing neurons.

## Discussion

Chicks with SBA-like features showed various neurological complications, including pain-related behavior (Fig. 1 and Suppl. Video 1) and motor dysfunction [21], consistent with reports of human neonates with SBA [6, 8, 10]. Therefore, pathological characterization of spinal motor networks of SBA chicks would provide insight into the progression of SBA-like neurological complications.

Spinal cord injury-induced pain complications are due in part to neuronal dysfunction in the spinal cord characterized by decreasing inhibitory controls [30–33]. In fact, loss of spinal cord dorsal horn inhibitory circuits, many of which involve interneurons that express GABA and its subpopulations, is a major contributor to persistent pain [13–18]. In this study, GABA- and PV-, CB-, and CR-expressing neurons were lost, and the expression levels of these factors were reduced in the exposed spinal cord dorsal horns of SBA chicks at the neonatal stage (Figs. 2–6). Therefore, reduction in GABAergic transmission in the spinal cord dorsal horn may have disrupted the inhibitory networks and contributed to the increased pain exhibited by SBA chicks. This was supported by the abnormal sensory projections in the dorsal funiculi of SBA chicks [20], which is the main route of inhibitory pain pathways in the spinal cord [34]. Although it is not possible to quantify pain in chicks, our behavioral observations suggested that the SBA chicks experienced persistent pain. The SBA chicks produced an increased number of loud vocalizations (Fig. 2b, c) and exhibited reduced confidence in mobility, escape reactions, anxiety, fear or restlessness, and struggling to change position (Suppl. video 1), major indicators of pain in birds [25, 26]. Thus, suppression of GABAergic transmission in the superficial dorsal horn may cause SBA-related pain.

Our findings also support a possible correlation between the loss of GABAergic neurons and neurological disorders in SBA patients, because intrathecal treatment with a GABA agonist relieves SBA-induced pain and spasticity in humans [35]. This also indicates that the reduction in spinal GABAergic transmission in the motor neuron area observed in our previous study contributed to leg movement dysfunction in the SBA chicks [21]. The loss of GABAergic transmission from the dorsal and ventral horn is also supported by the decreased or absent sensorimotor function in neonatal SBA chicks (Table 1). Taken together, these findings provide strong pathophysiological evidence that the loss of GABAergic transmission leads to varying degrees of motor and/or sensory deficits in the spinal cord, and that this contributed to various neurological complications, particularly leg movement dysfunction and pain, in SBA chicks.

Neuronal loss due to apoptosis is an important pathophysiological component of many neurological diseases [36–38]. The loss of spinal neurons at lesion sites may be associated with SBA-related neurological dysfunctions [39]. In fact, Kowitzk *et al*. suggested that neurons in neonatal myelomeningoceles undergo degeneration by apoptosis [40]. Apoptosis-induced degeneration of GABAergic interneurons and motor neurons in the lumbar cord ventral horn area is reportedly associated with SBA-like motor dysfunctions [21]. In this study, caspase-3 immunoreactivity, a marker of apoptosis, in GABAergic neurons in the dorsal horns of SBA chicks was evident at an advanced gestational stage (ED-18). Caspase-3 immunoreactivity gradually increased at the early neonatal stage and was strong at PD-10, when SBA chicks exhibited only a few damaged interneuron-like structures in the dorsal horn area (Fig. 7). Collectively, these findings demonstrate that reduced inhibitory transmission and increased apoptotic activity in the exposed spinal cord are involved in the pathogenesis of neurological complications, such as motor deficits and pain, in SBA chicks.

In conclusion, our findings provide evidence that early neonatal loss of inhibitory transmission disrupts the neuronal networks in the spinal cord dorsal horn, and that this contributes to SBA-related pain-like complications. Moreover, our results shed light on the mechanisms underlying cellular abnormalization and degeneration. They also suggest that the effects of neurological disorders in SBA patients can be modulated by manipulating the GABAergic system, which may be a promising avenue for novel therapeutic interventions.

## Acknowledgements

We would like to thank D. Shimizu for his technical support with confocal imaging.

## Author contributions

M.S.I.K. and S.M. planned, designed and performed experiments, analyzed results and compiled the manuscript; M.S.I.K. generated surgery-induced SBA chicks; H.N., F.I., T.S. S.S. and T.T. involved in data interpretation and manuscript editing; All authors discussed the study and approved the final manuscript.

## Additional Information

The authors declare no competing financial interests.

The English in this document has been checked by at least two professional editors, both native speakers of English. For a certificate, please see: http://www.textcheck.com/certificate/xzJOfn

---

**Supplementary Video 1.** Representative video clips of the behaviors of normal and SBA chicks at the early neonatal stage.

**Supplementary Video 2.** Representative video clips of the sensorimotor reactions to toe-pinching in normal and SBA chicks at the early neonatal stage.

